# Double-layer flexible neuronal probe with closely spaced electrodes for high-density *in-vivo* brain recordings

**DOI:** 10.1101/2020.10.15.341263

**Authors:** Sara Pimenta, José A. Rodrigues, Francisca Machado, João F. Ribeiro, Marino J. Maciel, Oleksandr Bondarchuk, Patrícia Monteiro, João Gaspar, José H. Correia, Luis Jacinto

**Affiliations:** CMEMS-UMinho, Department of Industrial Electronics, University of Minho, Guimarães, Portugal; Life and Health Sciences Research Institute (ICVS), School of Medicine, University of Minho, Braga, Portugal; ICVS/3B’s–PT Government Associate Laboratory, Braga/Guimarães, Portugal; International Iberian Nanotechnology Laboratory (INL), Braga, Portugal

## Abstract

Flexible probes for brain activity recordings are an attractive emerging approach that reduces mechanical mismatch between probe and neuronal tissue, thus minimizing the risk of brain damage or glial scaring. Although promising, flexible probes still present some technical challenges namely: i) how to overcome probe buckling during brain insertion given its intrinsically low mechanical rigidity; ii) how to fabricate closely spaced electrode configurations for high density recordings by standard lithography techniques in the flexible substrate. Here, we present a new flexible probe based solely on standard and low-cost lithography processes, which has closely spaced 10 μm diameter gold electrode sites on a polyimide substrate with inter-site distances of only 5 μm. By using a double-layer design and fabrication approach we were able to accommodate closely spaced electrode sites at two different depths from probe surface while also providing additional stiffening, just sufficient to prevent probe buckling during brain insertion. Detailed probe characterization through metrology of structural and electrical properties and chemical composition analysis, as well as functional assessment through *in vivo* high-density recordings of neuronal activity in the mouse cortex, confirmed the viability of this new fabrication approach and that this probe can be used for obtaining high quality brain recordings with excellent signal-to-noise ratio (SNR).

## Introduction

Silicon-based probes have taken the stage of neuronal recordings over the last decade, especially for acute applications. These probes can be fabricated through highly reproducible procedures, support many conductive electrodes and exhibit great stability ^1–3^. However, due to the intrinsic stiffness of the silicon substrate, there has been a rising interest in probes that can be fabricated in flexible biocompatible substrate materials with lower mechanical mismatch between probe and brain tissue ^4^. Flexible neuronal probes can be highly conformable, thus reducing the extent of brain tissue displacement, damage and glial scaring upon implantation, and providing adaption to brain micro- and macro-motions ^4,5,6,7^. These advantages are important for the stability of the brain implant and for improving the quality of the recorded signals, both in acute and chronic *in vivo* applications ^8^. Therefore, several flexible neuronal probes based on polyimide, SU-8 or Parylene C substrates have been recently proposed with various designs ^4,9^.

Nevertheless, integration of closely spaced electrodes for high density brain recordings, with minimal probe thickness and width, has not been widely sough-after in flexible probes due to added difficulties in integrating such designs with standard micromachining processes in flexible substrates. Electrode site size and density in neuronal probes is typically limited by the space needed to implement the vias that connect the electrode sites to the contact pads, while maintaining the size of the probe as small as possible. When considering standard lithography processes, although more affordable, they present lower resolution than for example electron- or ion-beam lithography, preventing patterning of sub-micron vias. On top of this, reducing the size of the electrodes or vias beyond a certain limit can increase electrode site impedance, and running vias too close to each other can increase parasitic capacitances, regardless of the fabrication method used. These factors can lead to signal distortion and low signal-to-noise (SNR) ratio thus adding to the challenge of fabricating closely spaced electrodes in flexible substrates through standard and low-cost lithography processes.

The desired flexibility of neuronal probes also creates challenges during brain insertion/implantation, since their intrinsically low mechanical rigidity may cause buckling. The unintended deformation of the probe at this stage can prevent brain penetration or result in misplacement at the target of interest ^4^. To overcome these limitations various mechanical augmentation strategies that increase probes’ buckling force threshold have been applied, namely coating the probes with absorbable molecules, such as polyethylene glycol (PEG) or saccharose, which are rigid at room temperature but dissolve when in prolonged contact with brain tissue; or the use of hard shuttle/carrier devices that help guiding the probe into the tissue and are then removed ^4,10^. But besides adding cumbersomeness to brain implantation procedures, these approaches typically increase the overall thickness of the inserted device, especially in the case of the shuttle which can be several microns larger than the probe itself. This leads to further tissue displacement causing additional brain damage upon insertion, thus defeating one of the main advantages of flexible probes.

Here, we present a polyimide-based flexible neuronal probe with 32 closely spaced gold electrode sites, fabricated with standard and low-cost lithography processes, that does not require additional stiffening aids for brain insertion. The implemented double layer architecture, where intermediate metallization and polyimide passivation layers allow the integration of closely spaced electrodes at two different depths from the probe’s surface, also provides sufficient additional intrinsic stiffening to prevent buckling. With this probe we were able to successfully record *in vivo* neuronal activity from closely spaced electrode sites without resorting to any stiffening procedure for brain penetration and probe insertion. Before *in vivo* recordings, we also fully characterized the viability of this new fabrication approach by metrology of structural and electrical properties as well as chemical composition, using scanning electron microscopy (SEM), energy-dispersive x-ray spectroscopy (EDS), x-ray photoelectron spectroscopy (XPS) and electrochemical impedance spectroscopy (EIS). Given that the proposed fabrication process employs only standard lithography techniques, multiple probe geometries with different arrangements of closely spaced electrode sites can be easily achieved at low-cost while maintaining structural properties.

## Results and discussion

### Fabricated flexible probe

Figure 1 shows the design of the double-layer polyimide flexible probe with closely spaced electrode sites. The neuronal probe contains 32 circular gold electrode sites, each one with approximately 10 μm diameter. The inter-site distance is 5 μm in both directions, creating a closely spaced matrix of electrode sites. The neuronal probe has approximately 150 μm width, 24 μm thickness and 6 mm shaft length. In order to fabricate a high-density probe with closely spaced electrode sites in a flexible substrate, a two-layer design with electrode sites at two different depths (top and bottom) from the probe surface (6 and 12 μm, respectively) was implemented. Electrodes, vias, and contacts were distributed across two different depths and patterned in sequential deposition, photolithographic and reactive ion etching (RIE) steps. This strategy allows the electrode sites to be in close proximity and still have enough space for running the vias to the contact pads along the two layers, while keeping the probe with a small physical footprint. Polyimide was chosen as the substrate due to its superior stability, biocompatibility and low elastic modulus ^4,11^. By using a polyimide film with a Young’s elastic modulus of only 8.5 GPa as the substrate, it was ensured that the probe was flexible (Fig. 1b) and with a stiffness of at least an order of 10 lower than a silicon probe with the same dimensions ^12^. Additionally, the use of non-photodefinable polyimide permitted the implementation of low-cost etching fabrication procedures. Details of the fabrication process flow can be found in the Materials and Methods section.

**Figure 1.**
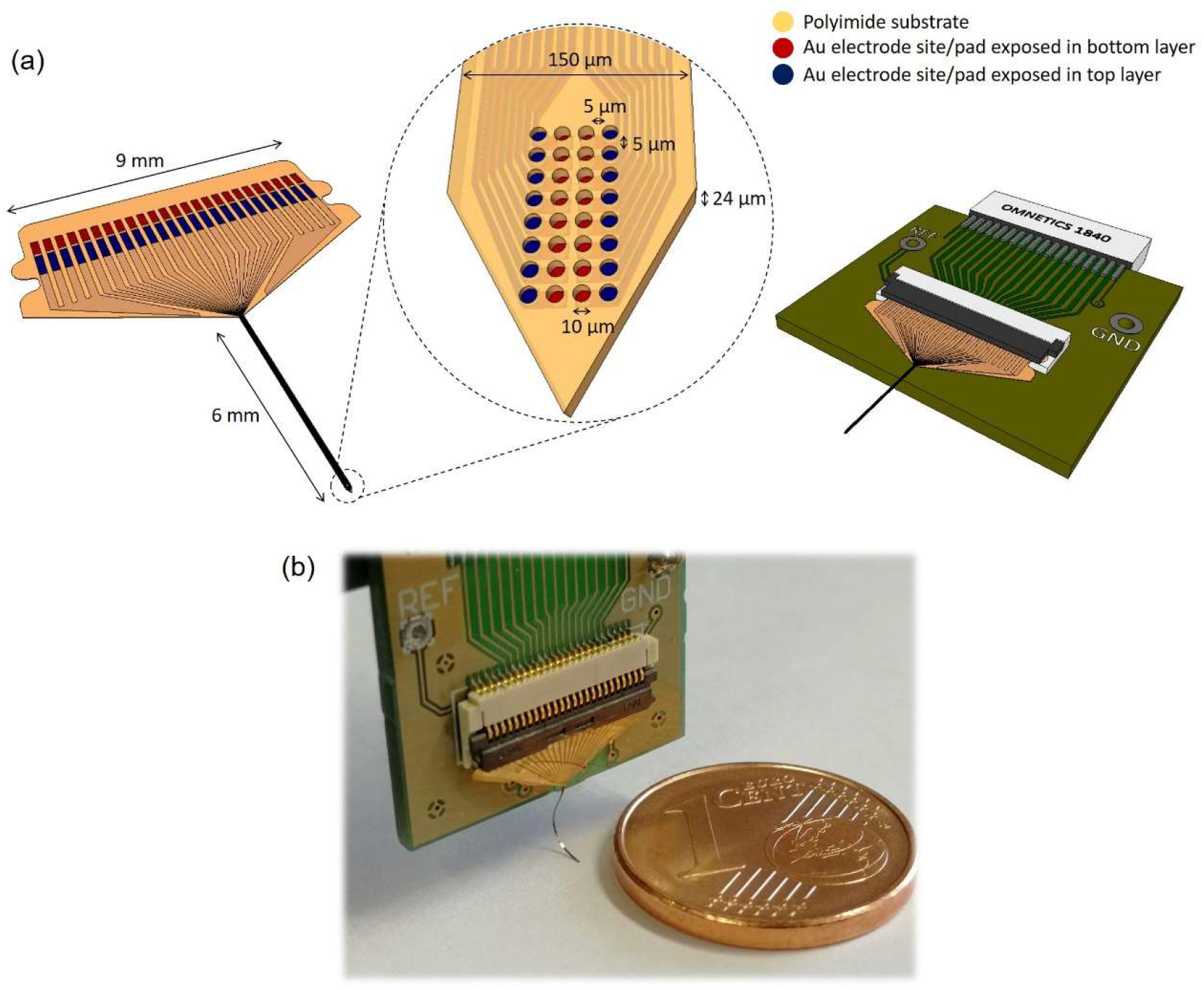
(a) Schematic illustration of the flexible polyimide neuronal probe design (left) and the printed circuit board (PCB) for packaging (right). The central inset shows the distribution of the closely spaced electrode sites at the tip of the probe. (b) Fabricated flexible neuronal probe coupled to the PCB.

Figure 2 shows SEM images of a fabricated probe with the exposed gold surface of the electrode sites at two different depths. The observed clear color difference between top and bottom electrode sites due to their different depths reveals correct polyimide passivation (Fig. 2a and 2b). The use of direct laser writing (DLW) to pattern the second metal layer allowed a higher resolution and thus improved the alignment between the top and bottom electrode sites. Figure 2c shows the bottom and top metal tracks patterned along the polyimide substrate and the absence of short circuit between these two metallic layers, separated by an intermediate polyimide layer.

**Figure 2.**
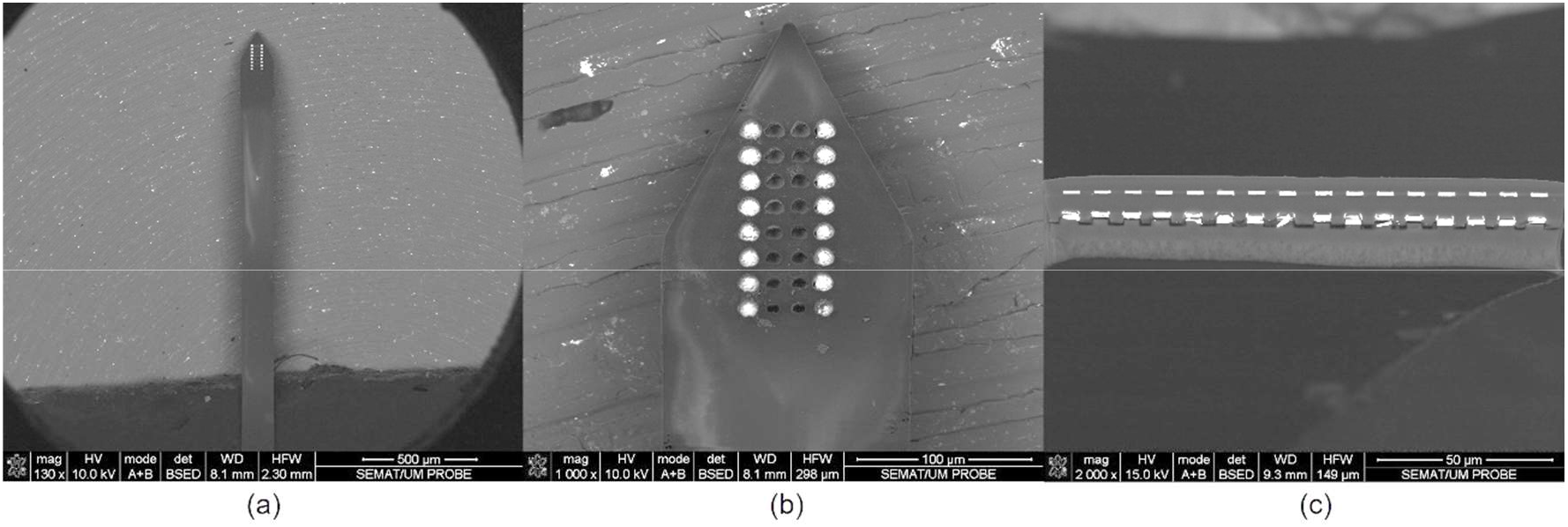
SEM images of the flexible polyimide neuronal probe: (a) top view of probe shaft and tip containing the electrode sites; (b) detail view of the probe tip, showing the closely spaced electrode sites at two different depths; and (c) cross-section view of the probe shaft showing two metallic vias separated by the intermediate polyimide layer.

### Chemical and electrical characterization of the fabricated flexible probe

To assess the viability of the new fabrication process, metrology of electrical properties and chemical composition analysis were performed by EDS, XPS and EIS.

Figure 3 shows the EDS and XPS results. EDS analysis revealed gold (Au) as the most prevalent element at the bottom and top electrode sites with a percentage by weight higher than 56 wt%, confirming successful etching during fabrication. Other detected elements include carbon (C) (higher than 24 wt%), tungsten (W) (higher than 3 wt%) and aluminum (Al), titanium (Ti), oxygen (O), copper (Cu) and Cerium (Ce) (all below 3 wt%). The presence of other elements in lower concentrations is expected, as C is in the polyimide layers, Al, Ti, Cu and W are part of the metal stacks and Ce and O are residuals of the wet and polyimide etching processes. The EDS analysis detected the chemical element C as being the most prevalent in the probe substrate (approximately 80 wt%), as expected from the presence of a carbonyl group in the polyimide polymer film. XPS measurements further validated the presence of Au at the electrode sites surface. This analysis showed that the atomic concentration of Au at the probe surface increased as a function of etching time, while Ce decreased, further confirming successful etching processes during the fabrication of the flexible probe.

**Figure 3.**
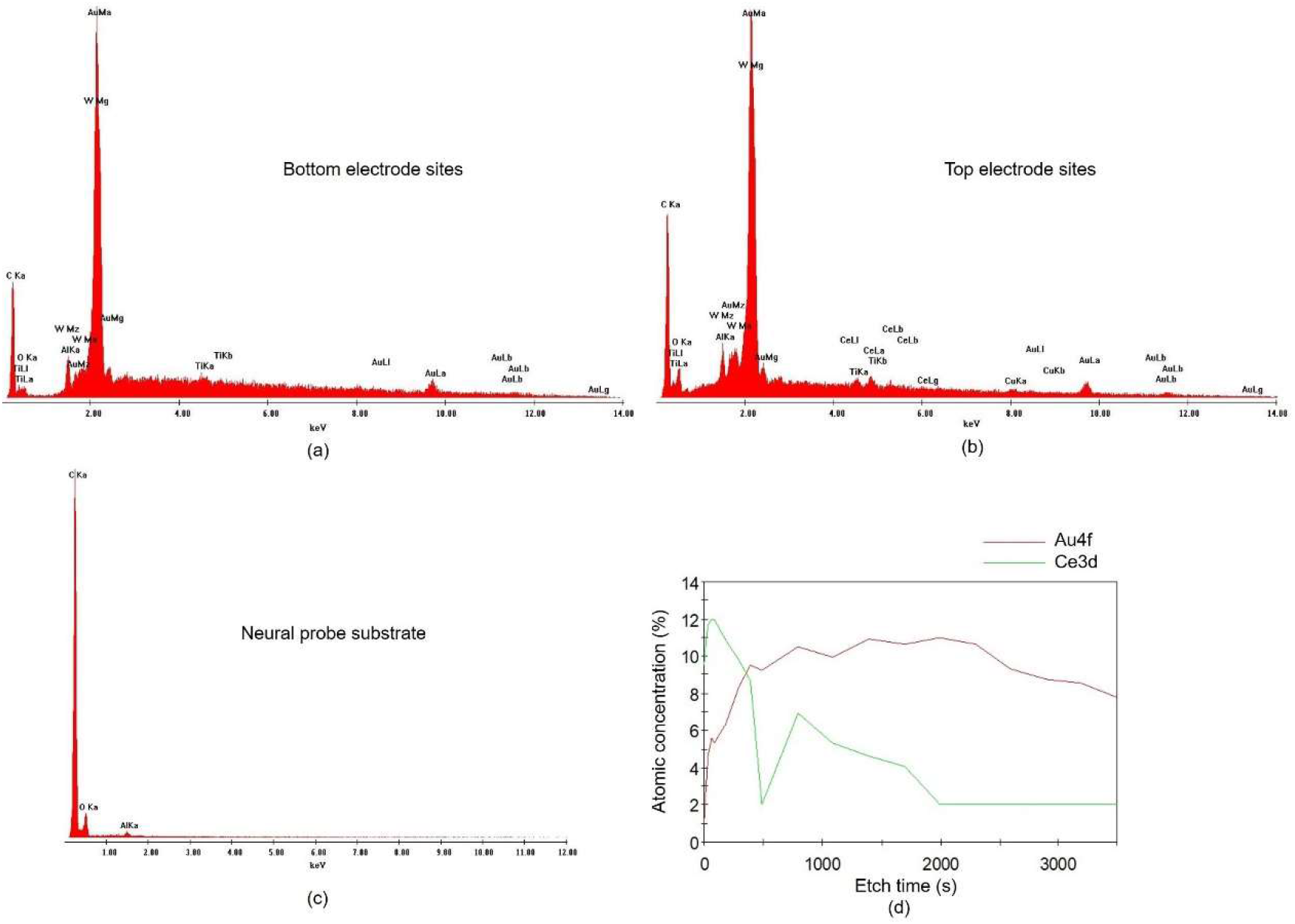
EDS and XPS results of the flexible polyimide neuronal probe: (a,b) chemical composition of a bottom (a) and a top (b) electrode sites with clear peaks for gold (Au); (c) chemical composition of the neuronal probe substrate with a clear peak for carbon (C); (d) XPS results for the probe surface showing an increase of gold (Au) as function of the etching time.

Following cleaning of the electrode sites with a potassium hydroxide (KOH) and hydrogen peroxide (H_2_O_2_) solution to obtain clean gold surfaces ^13^, EIS measurements were performed. The mean impedance magnitude of the 32 electrode sites at 1 kHz was approximately 100 kΩ. The fabrication of gold electrode sites with 10 μm diameter allowed electrodes with an impedance within the optimal range for neuronal recordings with low intrinsic noise and high signal-to-noise ratio (SNR) ^14,15^, without the need for any additional electrodeposition steps.

### In-vivo Electrophysiology

To perform functional assessment of the closely spaced electrode sites in the probe, electrophysiological brain recordings were performed *in vivo*. Figure 4 shows spontaneous and opto-evoked neuronal activity recorded from the motor cortex of anesthetized mice.

**Figure 4.**
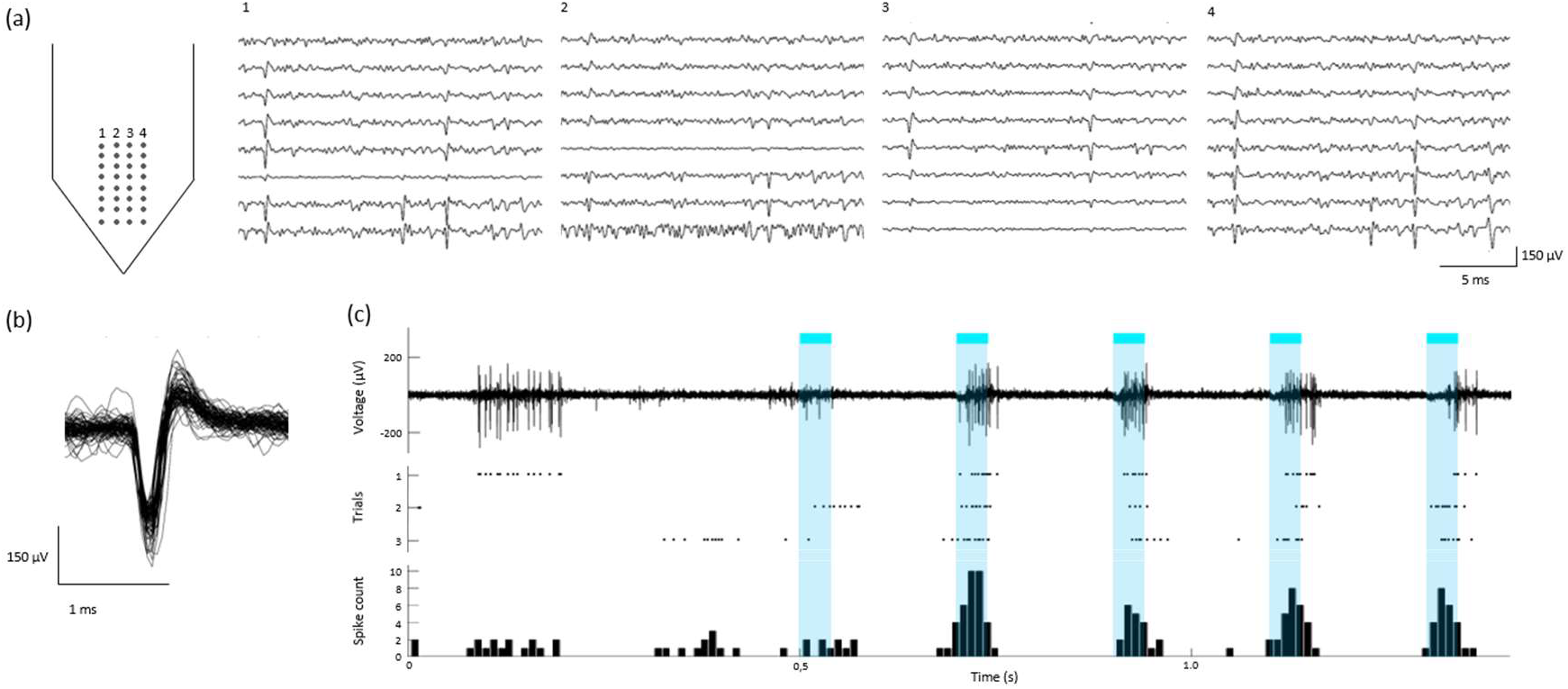
*In vivo* recordings in mice cortex: (a) example of neuronal activity recorded simultaneously from all channels of the probe (signals band-pass filtered between 0.3 and 6 kHz); (b) example of an isolated single-unit from the same recording, showing the first 100 spike waveforms; (c) Opto-evoked neuronal activity with an example of signal trace (top; filtered 0.3 – 6 kHz), spike raster plot for 3 trials (middle) and cumulative peri-stimulus time histogram (PSTH) with total spike count across time for the 3 trials.

After removing dura and exposing the brain surface, it was possible to easily insert our flexible probe without buckling not having to resort to any insertion aid, contrarily to other flexible neuronal probes previously published ^4^. Although our probe is up to 10 μm thicker than some of these probes ^4^, it has a higher electrode site count per area as well as smaller inter-site distance, while maintaining a low mechanical rigidity within the desired range for a flexible substrate probe.

Figure 4a shows an example of recorded neuronal activity across all channels in the probe, where it is possible to observe multiple spiking units across several recording sites. The average RMS from the signal across all channels was approximately 11 μV, with isolated spikes crossing a threshold of at least 5 times that value. The closely spaced configuration design allowed the recording of the same neuronal activity from the same neurons across several electrode sites. Although the electrode sites are at two different depths from the surface of our probe, correlated neuronal activity in the cortex at was reliably recorded at both depths and a conventional spike sorting algorithm was used to successfully isolate several single units from each recording. Figure 4b shows an example of an isolated single unit. It has been previously shown that spike sorting can be facilitated by spatial over-representation of neuronal activity across several sites ^14^, especially in recording devices where the electrode geometry is fixed, as observed here.

To further assess the capability of the probe in detecting modulated neuronal activity and performance in a scenario of an optogenetics experiment, a typical application sought after by many experimental neuroscience labs, neuronal responses were evoked from cortical neurons expressing channelrhodopsin by shining blue light into the cortical region. Figure 4c shows successful brain recordings, with excellent signal-to-noise ratio (SNR), of both spontaneous and opto-evoked neuronal activity.

## Conclusion

This paper presents a new flexible polyimide probe with closely spaced electrode sites for high-density neuronal recordings, which requires no additional coating/stiffening procedures to allow brain insertion. With the microfabrication process described here, it was possible to use standard and low-cost photolithography techniques to fabricate a double-layer high-density probe in a flexible substrate with low stiffness. The intermediate metal and polyimide passivation layers of the double layer design allowed a matrix arrangement of closely spaced electrodes while providing just sufficient additional stiffening to prevent buckling during brain insertion. The obtained size and mean impedance of the gold electrode sites (approximately 100 kΩ) facilitated *in vivo* recordings with high SNR and subsequent single-unit spike-sorting. Our results show that the probe presented here can be amenable for widespread use by removing the necessity of insertion aids and providing high SNR neuronal activity recordings. Additionally, since only standard and low-cost lithography processes were used it can be scaled to include, for example, multiple-shank designs and different electrode configurations on each shank just by simple changes to mask patterning.

## Materials and Methods

### Probe Fabrication

A standard photolithography process was employed, in which alternating polyimide and metallization layers were deployed, each followed by successive stages of RIE to pattern the electrode sites, vias and contact pads. Figure 5 shows a graphical depiction of the probe fabrication steps.

**Figure 5.**
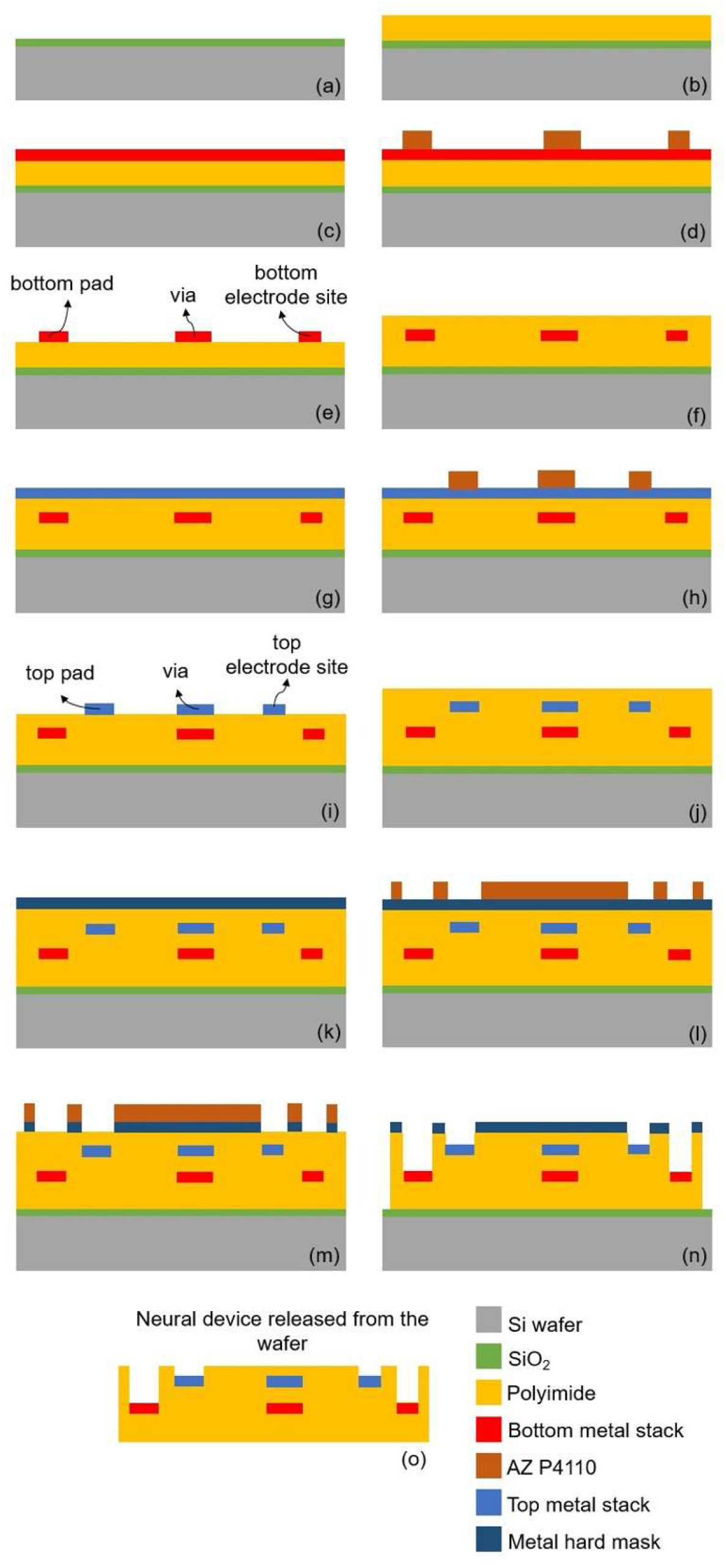
probe fabrication process (not to scale).

A sacrificial film (0.5 μm thick SiO_2_) was initially deposited by chemical vapor deposition (CVD) over a 200 mm silicon wafer to allow detachment of the probe at the end of processing (Fig. 5a). This was followed by spin-coating, soft-baking and curing a layer of non-photodefineable polyimide (PI-2611 from HD MicroSystems), approximately 7.5 μm thick, which formed the base of the probe (Fig. 5b). Then, a metal stack (TiW/AlSiCu/TiW (50/500/50 nm), Au (200 nm), Cr (10 nm) and AlSiCu/TiW (250/50 nm)) was sequentially sputtered (Fig. 5c). The electrode sites, vias and contact pads of what would become the bottom layer were then defined by UV photolithography with a Cr hard mask and photoresist (AZ P4110 from MicroChemicals), and patterned by RIE, followed by O_2_ stripping to remove the remaining photoresist (Fig. 5d and 5e). A second 7.5 μm thick layer of PI-2611 was again spin-coated, baked and cured to ensure passivation of the bottom metal layer (Fig. 5f). A second metal stack (TiW/AlSiCu/TiW (50/500/50 nm), Au (200 nm), Cr (10 nm) and AlSiCu/TiW (350/50 nm)) was sputtered sequentially (Fig. 5g). The electrode sites, vias and contact pads of the top layer were defined by UV photolithography with direct laser writing laser (DLW) and photoresist (AZ P4110 from MicroChemicals), and patterned by RIE, followed by O_2_ stripping to remove the remaining photoresist (Fig. 5h and 5i). A final layer of PI-2611 was spin-coated, soft-baking and cured to ensure passivation of the probe (Fig. 5j). A metal hard mask (AlSiCu/TiW (600/50 nm)) was sputtered (Fig. 5k), defined by UV photolithography with a Cr hard mask and photoresist (AZ P4110 from MicroChemicals), and patterned with RIE (Fig. 5l and 5m). Then, an O_2_/CF_4_ etching process was performed for PI-2611 etching, exposing the electrode sites and contact pads of both the bottom and top electrode layers (Fig. 5n). Finally, two wet etching processes (AlSiCu and Cr etching) and an HF vapor etching process (SiO_2_ etching) were performed to remove the hard mask, to expose the Au in the electrode sites and contact pads and to release the neuronal probe from the silicon wafer (Fig. 5o).

### Probe Packaging

The released probes were packaged with a custom-designed printed circuit board (PCB) to allow access to the electrodes through an headstage (RHD2132, Intan), which includes a pre-amplifier and a digitizer, using an omnetics connector (A79022-001). An FPC connector (FH29DJ-50S-0.2SHW from Hirose Electric) connected the flexible contact pads on the probe side to the PCB. The PCB package allowed the electrode site gold surface cleaning procedure, signal acquisition for impedance measurements and *in vivo* recordings.

In order to obtain clean gold surfaces at the electrode sites, removing unwanted residuals from the fabrication process, the tip containing the electrode sites was immersed in a solution of 50 mM potassium hydroxide (KOH) and 25% of hydrogen peroxide (H_2_O_2_) for 10min, as described in ^13^, followed by a 50 mM KOH sweep, using a potentiostat (Reference 600, Gamry Instruments) and a sweep from −200 to −1200 mV (vs. Ag/AgCl) at 50 mV/s scan rate.

### Probe characterization

SEM was performed to structurally evaluate the fabricated probe. SEM measurements were performed at SEMAT/University of Minho, using a NanoSEM - FEI Nova 200 equipment.

EDS was performed to chemically evaluate the probe and confirm that the electrode sites were etched correctly to expose the gold surface. EDS was performed during SEM measurements, also at SEMAT/University of Minho, using an EDAX - Pegasus X4M equipment. XPS was also performed to chemically evaluate the surface of the probe at a depth of approximately 10 nm. XPS measurements of Au and Ce atomic concentrations were performed using a Thermo Scientific Escalab 250 Xi equipment, and considering an analysis area of 650 μm diameter, as a function of the etching time.

Electrochemical impedance spectroscopy (EIS) measurements were performed with a potentiostat (Reference 600, Gamry Instruments) in a three electrode configuration in 0.9% saline electrolyte: working electrode (each electrode site of the probe), counter electrode (Pt wire) and reference electrode (Ag/AgCl). An AC sinusoidal signal of 5 mV with a frequency between 100 Hz and 10 kHz and was applied, and the impedance value at 1 kHz for each microelectrode was recorded.

### Electrophysiology and Optogenetics

Emx1-Cre;Ai27D male mice originally purchase from JAX (stocks #005628, #012567) ^16,17^ were used in this study. All experiments were conducted in accordance with European Union Directive 2016/63/EU and the Portuguese regulations and laws on the protection of animals used for scientific purposes (DL Nº 113/2013). This study was approved by the Ethics Subcommittee for Life and Health Sciences of University of Minho (ID: SECVS 01/18) and the Portuguese Veterinary General Direction (ID: DGAV 8519).

Briefly, mice were anesthetized by an intraperitoneal injection of ketamine (75 mg/kg) and medetomidine (1 mg/kg) mix and positioned in a stereotaxic frame (Stoelting). Following a midline incision across the skull, a burr hole above the motor cortex (M1) (1.1mm AP and 1.5mm ML) ^18^ exposed the surface of the brain. After careful removal of the dura, the probe, connected to the PCB and attached to a precision micrometric stereotaxic arm (1760, Kopf Instruments), was lowered to at least 250 μm from the brain surface. A stainless-steel screw, in another burr hole at the back of the skull, served as ground. Extracellular signals were acquired with an Open Ephys acquisition system at 30 kS/s.

To evoke neuronal activity from M1 neurons expressing light-gated channelrhodopsin, the surface of the exposed brain was illuminated with light pulses (473 nm wavelength, 5 Hz with, 50 ms pulse width) by an optical fiber (200 μm diameter), held by a micromanipulator, and connected to a DPSS laser source (CNI). Pulse triggering was delivered by a waveform generator (DG1022, Rigol) connected to the laser source.

Signals were analyzed with custom-writen Matlab code (Mathworks) and JRClust ^19^. Extracellular recordings were filtered between 0.3 and 6 kHz and spikes were detected using an amplitude threshold at least 5 times higher than an estimate of background noise standard deviation ^20^. Initial spike sorting, where spikes were assigned to different unit clusters, was automatically performed by JRClust, followed by manual cluster curation based on visual inspection of spike waveforms, inter-spike interval histograms and auto-/cross-correlograms.

## Acknowledgements

This work was supported by FCT with: project OpticalBrain reference PTDC/CTM-REF/28406/2017, operation code POCI-01-0145-FEDER-028406, through the COMPETE 2020; CMEMS-UMinho Strategic Project UIDB/04436/2020; and Infrastructures Micro&NanoFabs@PT, operation code NORTE 01-0145-FEDER-022090, POR Norte, Portugal 2020. This work has also been funded by National funds, through the Foundation for Science and Technology (FCT) - project UIDB/50026/2020 and UIDP/50026/2020; the projects NORTE-01-0145-FEDER-000013 and NORTE-01-0145-FEDER-000023, supported by Norte Portugal Regional Operational Programme (NORTE 2020), under the PORTUGAL 2020 Partnership Agreement, through the European Regional Development Fund (ERDF); FCT project PTDC/MED-NEU/28073/2017 (POCI-01-0145-FEDER-028073); and by The Branco Weiss fellowship - Society in Science, (ETH Zurich).

## Contributions

The work presented in this paper was a collaboration of all authors. Sara Pimenta, João F. Ribeiro, João Gaspar and José H. Correia conceived and designed the flexible probe. The fabrication process was performed by Sara Pimenta and João F. Ribeiro and supervised by João Gaspar. The probe characterization was performed by José A. Rodrigues, Marino J. Maciel and Oleksandr Bondarchuk. José. A. Rodrigues and Sara Pimenta analyzed the probe characterization results. Luis Jacinto designed and performed the *in vivo* experiments. Patricia Monteiro provided the transgenic animals and performed the optogenetic experiments with Luis Jacinto. Francisca Machado analyzed the neuronal activity data. The first draft of the manuscript was written by Sara Pimenta and José A. Rodrigues. José H. Correia and Luis Jacinto revised the manuscript. José H. Correia and Luis Jacinto supervised all the fabrication and animal validation works, respectively.

## References

1. Buzsáki, G. et al. Tools for probing local circuits: High-density silicon probes combined with optogenetics. Neuron (2015) doi:10.1016/j.neuron.2015.01.028.

2. Jun, J. J. et al. Fully integrated silicon probes for high-density recording of neural activity. Nature (2017) doi:10.1038/nature24636.

3. Ulyanova, A. V. et al. Multichannel silicon probes for awake hippocampal recordings in large animals. Front. Neurosci. (2019) doi:10.3389/fnins.2019.00397.

4. Weltman, A., Yoo, J. & Meng, E. Flexible, penetrating brain probes enabled by advances in polymer microfabrication. Micromachines (2016) doi:10.3390/mi7100180.

5. Seymour, J. P., Wu, F., Wise, K. D. & Yoon, E. State-of-the-art MEMS and microsystem tools for brain research. Microsystems Nanoeng. 3, 16066 (2017).

6. Spencer, K. C., Sy, J. C., Falcón-Banchs, R. & Cima, M. J. A three dimensional in vitro glial scar model to investigate the local strain effects from micromotion around neural implants. Lab Chip (2017) doi:10.1039/c6lc01411a.

7. Moshayedi, P. et al. The relationship between glial cell mechanosensitivity and foreign body reactions in the central nervous system. Biomaterials (2014) doi:10.1016/j.biomaterials.2014.01.038.

8. Kozai, T. D. Y., Jaquins-Gerstl, A. S., Vazquez, A. L., Michael, A. C. & Cui, X. T. Brain tissue responses to neural implants impact signal sensitivity and intervention strategies. ACS Chem. Neurosci. (2015) doi:10.1021/cn500256e.

9. Fan, B. et al. Flexible, diamond-based microelectrodes fabricated using the diamond growth side for neural sensing. Microsystems Nanoeng. 6, 42 (2020).

10. Na, K. et al. Novel diamond shuttle to deliver flexible neural probe with reduced tissue compression. Microsystems Nanoeng. 6, 37 (2020).

11. Rubehn, B. & Stieglitz, T. In vitro evaluation of the long-term stability of polyimide as a material for neural implants. Biomaterials 31, 3449–3458 (2010).

12. Lecomte, A., Descamps, E. & Bergaud, C. A review on mechanical considerations for chronically-implanted neural probes. J. Neural Eng. 15, 031001 (2018).

13. Fischer, L. M. et al. Gold cleaning methods for electrochemical detection applications. Microelectron. Eng. 86, 1282–1285 (2009).

14. Scholvin, J. et al. Close-packed silicon microelectrodes for scalable spatially oversampled neural recording. IEEE Trans. Biomed. Eng. (2016) doi:10.1109/TBME.2015.2406113.

15. Neto, J. P. et al. Does Impedance Matter When Recording Spikes With Polytrodes? Front. Neurosci. 12, (2018).

16. Gorski, J. A. et al. Cortical excitatory neurons and glia, but not GABAergic neurons, are produced in the Emx1-expressing lineage. J. Neurosci. 22, 6309–6314 (2002).

17. Madisen, L. et al. A toolbox of Cre-dependent optogenetic transgenic mice for light-induced activation and silencing. Nat. Neurosci. 15, 793–802 (2012).

18. Franklin, K. & Paxinos, G. The Mouse Brain in Stereotaxic Coordinates. (Academic Press, 2008).

19. Jun, J. et al. Real-time spike sorting platform for high-density extracellular probes with ground-truth validation and drift correction. bioRxiv (2017) doi:10.1101/101030.

20. Quiroga, R. Q., Nadasdy, Z. & Ben-Shaul, Y. Unsupervised spike detection and sorting with wavelets and superparamagnetic clustering. Neural Comput. 16, 1661–1687 (2004).

